# The steroid-inducible pOp6/LhGR gene expression system is fast, sensitive and does NOT cause plant growth defects in rice (*Oryza sativa*)

**DOI:** 10.1101/2021.01.02.425069

**Authors:** Marketa Samalova, Ian Moore

## Abstract

Inducible systems for transgene expression activated by a chemical inducer or an inducer of non-plant origin are desirable tools for both basic plant research and biotechnology. Although, the technology has been widely exploited in model plants, it has not been optimised for use with the major monocotyledonous crop species, namely rice. We have adapted the dexamethasone-inducible pOp6/LhGR system for rice and shown that it is fast, sensitive and tightly regulated, with high levels of induction that remain stable over several generations. Most importantly, we have shown that the system does not cause negative growth defects *in vitro* or in soil grown plants. Interestingly in the process of testing, we found that another steroid, triamcinolone acetonide, is a more potent inducer in rice than dexamethasone. We present serious considerations for the construct design to avoid undesirable effects caused by the system in plants, leakiness and possible silencing, as well as simple steps how to maximize translation efficiency of a gene of interest. Finally, we compare the performance of the pOp6/LhGR system with other chemically inducible systems tested in rice in terms of the properties of an ideal inducible system.

**Significance statement:** The non-monocot codon-optimized version of the dexamethasone inducible pOp6/LhGR system does not cause severe developmental perturbations in rice plants.

## INTRODUCTION

Chemically inducible systems that regulate gene expression are crucial tools for basic plant biology research as well as biotechnology applications. They allow analysis of gene primary effects before homeostatic mechanisms start to counteract and reveal a clear correlation between induction of a transgene and occurrence of an altered phenotype. Their applications include expression of gene products that interfere with regeneration, growth or reproduction; regulation and expression at different stages of plant development; conditional genetic complementation, co-suppression and overexpression studies. However, adaption of existing inducible systems developed in model plants to other species including monocots and important crop plants particularly rice, has not been easily achieved.

The systems typically contain two transcription units. The first unit employs a constitutive or tissue-specific promoter to express a chemical-responsive transcription factor, the second unit consists of multiple copies of the transcription factor binding site linked to a minimal plant promoter, which is used to express the target gene. The most optimal systems developed to date include dexamethasone-inducible GVG (Aoyama and Chua, 1997) and pOp6/LhGR (Craft *et al*., 2005; Samalova *et al*., 2005), estrogen-inducible XVE (Zuo *et al*., 2000), ecdysone agonist-inducible VGE (Padidam *et al*., 2003; Koo *et al*., 2004) and ethanol-inducible *alc* systems (Caddick *et al*., 1998; Roslan *et al*., 2001; Salter *et al*., 1998). Their characteristics were compared in detail by Moore *et al*., (2006) and potential applications in plant biotechnology were reviewed by Corrado and Karali, (2009).

Several attempts have been made to develop an inducible system for gene manipulation in rice. The XVE system was tested by Sreekala *et al*., (2005) and Okuzaki *et al*., (2011) but the use remains very limited due to the lack of systemic movement of the estradiol inducer (Hirose *et al*., 2012, Chen *et al*., 2017). Two other steroid-inducible systems have been tested but the GVG and a modular gene expression system (Vlad *et al*., 2019) derived from the pOp6/LhGR system met with a mixed success in rice, as they caused severe growth and developmental defects of the plants (Ouwerkerk *et al*., 2001; Vlad *et al*., 2019).

An easier and more common strategy exploited in rice is the use of conditional promotors that can be activated by heat (Khattri *et al*., 2011; Sun *et al*., 2012; Rerksiri *et al*., 2013), pathogens (Helliwell *et al*., 2013; Quilis *et al*., 2014) or wounding (Quilis *et al*., 2014), oxidative (Woo *et al*., 2015) and other stresses (Nakashirma *et al*., 2007). Other crops promoters include induction by heat in maize (Du *et al*., 2019) and potato (Kopertekh *et al*., 2018), by light in tomato (Timerbaev and Dolgov, 2019) or, cold in barely (Eva *et al*., 2018), wheat (Meszaros *et al*., 2015) and sweet potato (Honma *et al*., 2019). Often the conditional expression is combined with the Cre-lox technology (Khattri *et a*l., 2011; Meszaros *et al*., 2015; Eva *et al*., 2018; Kopertekh *et al*., 2018; Du *et al*., 2019) or an alternative site-specific recombinase system (Woo *et al*., 2015) to generate marker-free genetically modified plants.

Adaptation of the breakthrough CRISPR/Cas9 technology to plants (Jiang *et al*., 2013) including rice (Zhang *et al*., 2014) allows genome editing and creates a powerful tool for engineering knockdown, knockout or chimeric plants. This genome editing system was combined with a heat-shock-inducible promoter to generate heritable mutations in rice (Nandy *et al*., 2019) and a virus-inducible system was developed that confers resistance to Gemini viruses in model plants *Arabidopsis* and tobacco (Ji *et al*., 2018). Recently, the technology was integrated with the estradiol-inducible XVE-based cell-type-specific system (Siligato *et al*., 2016; Zuo *et al*., 2000) to create an inducible genome editing (IGE) system in *Arabidopsis* that enables efficient generation of target gene knockouts in desired cell types at different developmental stages (Wang *et al*., 2020).

Over the years we and others have made considerable efforts to develop the dexamethasone-inducible transcription activation system, pOp6/LhGR, as a tool for the growing demands on modern gene technologies. It is a widely used system for which a comprehensive library of cell-type specific activator lines was created in *Arabidopsis* (Schurholz *et al*., 2018); the system was combined with artificial micro-RNA (amiRNA) to knockdown multigene expression (Goth *et al*., 2012; Samalova *et al*., 2020) and hairpin RNAi molecules to silence gene expression (Wielopolska *et al*., 2005; Liu and Yoder, 2016). Apart from *Arabidopis* (Craft *et al*., 2005) and tobacco (Samalova *et al*., 2005) the system was tested in various other species including citrus plant (Rossignol *et al*., 2014) and *Medicago truncatula* (Liu and Yoder, 2016) and most recently in rice (Vlad *et al*., 2019). However, the specific modifications made to the original version of the pOp6/LhGR system caused undesirable effects in plants.

This report describes the functionality of the pOp6/LhGR system in rice, its stability over several generations, time course and dose response characteristics; optimization of induction by various steroids as inducers and methods of systemic and localised applications that do not have any detrimental effects in rice even after prolonged induction.

## RESULTS

### Evidence that the pOp6/LhGR system is functional in rice

The pOp6/LhGR system (Craft *et al*., 2005; Samalova *et al*., 2005) comprises of a transcription activator LhGR which is a fusion between a high-affinity DNA-binding mutant of *Escherichia coli lac* repressor, *lacI*^His17^, transcription-activation-domain-II of Gal4 from *Saccharomyces cerevisiae* and the ligand-binding domain (LBD) of the rat glucocorticoid receptor (GR). The pOp6 is a chimeric promoter that consists of six copies of *lac* operators (*lacOp*) cloned upstream of a minimal cauliflower mosaic virus (*CaMV*) 35S promoter (−50 to +8) and is apparently silent when introduced into plants. The principle of the system is that in the absence of the steroid ligand, dexamethasone (Dex), the transcription factor is trapped in an inactive complex via interaction of the GR LBD and heat-shock protein HSP90. Upon induction with Dex, this complex is disrupted and the LhGR activator binds to the pOp6 promotor and induces expression of the target gene of interest.

To adopt the pOp6/LhGR system for rice, first we chose a binary pVec8-overexpression vector (Kim and Dolan, 2016) in which we placed the LhGR2 activator sequence that incorporates the *Arabidopsis* codon-optimized *GAL4* sequence (Rutherford *et al*., 2005) under the control of a maize ubiquitin promoter that contains an intron (*pZmUbi*). The inclusion of an intron is well known to greatly increase expression efficiency in monocots, but similar effects have been reported in dicots and other eukaryotes (Rose, 2004). Secondly, we checked in the literature (Mitsuhara *et al*., 1996; Segal *et al*., 2003) that none of the sequences of the pOpIn2 bidirectional reporter cassette (Samalova *et al*., 2019) including the *lacOp*, minimal promoters and tobacco mosaic virus (*TMV*) omega (Ω) translation enhancers have been reported to be toxic or non-functional in monocots and cloned it into the activator construct to create pVecLhGR2 as depicted in Fig. 1 and Supp. Fig. 1. For simplicity of testing the regulated expression of the pOp6/LhGR in rice, we used the *uidA* (encoding β-glucuronidase; GUS) and the yellow fluorescent protein (*YFP*) as the genes of interest.

**Figure 1:**
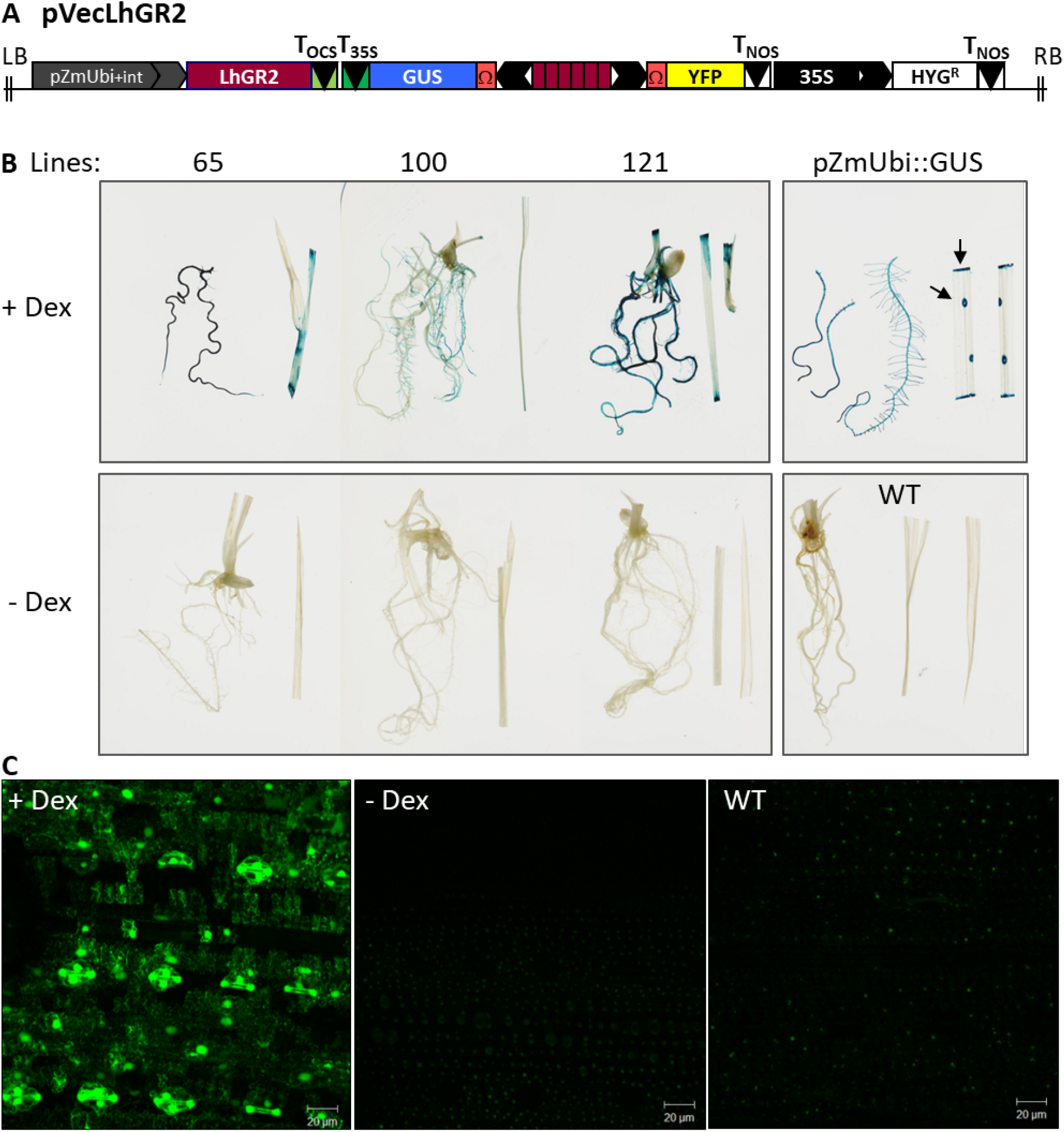
The pOp6/LhGR system in rice: a proof of principle. **(A)** A schematic representation of the pVecLhGR2 construct that contains the *Arabidopsis* codon-optimized *GAL4* sequence (Rutherford *et al*., 2005) of the transcriptional activator LhGR2 driven by a *pZmUbi* promoter (containing an intron), a bidirectional pOp6 promotor version with two *TMV* Ω translation enhancers driving two reporter genes -*uidA* (encoding β-glucuronidase; GUS) and a yellow fluorescence protein (*YFP*), and a pCaMV35S::HYG selectable marker cassette conferring hygromycin resistance. The construct was assembled as detailed in Supplementary Figure S1. **(B)** Twelve-day-old rice seedlings of three independent transgenic lines (number 65, 100 and 121) histochemically stained for GUS activity that was induced by seedling transfer onto ½ MS plates containing 30 μM Dex or control DMSO (-Dex) for 6 days. Seedlings of pZmUbi::GUS and wild-type (WT) were included as controls. The arrows point to an example of damaged cells/ cuts where the GUS staining substrate (5-bromo-4-chloro-3-indolyl glucuronide; X-Gluc) penetrated inside the cells. **(C)** Leaves of 4-week-old plant (line 121) were painted with 30 μM Dex or control solution (-Dex) and imaged using a confocal laser scanning microscope 96 h later to detect YFP fluorescence. WT was included as a control for autofluorescence. The scale bar is 2 μm.

To create stable transgenic rice lines, we used a protocol for *Agrobacterium-*mediated transformation of calli induced from seeds of *Oryza sativa spp. japonica* cultivar Kitaake as described by Vlad *et al*., 2019. We generated several independent transgenic lines (T1) in which the induced GUS staining was comparable to that from the constitutive promoter *pZmUbi* (Fig. 1B) and the *YFP* expression (Fig. 1C) was inducible by Dex, proving that the pOp6/LhGR system is in principle functional in rice. However, the transformation efficiency was relatively low and to induce higher levels of expression various alternate methods were tested.

### Transformation efficiency and reliability of induction in subsequent generations

We tested GUS activity by histochemical staining in 120 generated putative transformants that included little plantlets with a piece of callus and some roots (Table 1). We induced them in liquid ½ MS with 30 μM Dex and stained for GUS activity 24h later. Approximately a half of them showed visible (by eye) GUS staining in roots after 2h and shoots after 4h. The reaction was stopped and scored at 24h with the following pattern: 30.8% root staining only, 9.2% shoots only, 12.5% stained shoots, roots and callus (data not shown).

**Table 1:** Number of putative transformants and generated transgenic lines tested for hygromycin (HYG) resistance and induction of GUS activity in subsequent generations.

| Generation | No. plants tested | HYG resistant | GUS staining |
| --- | --- | --- | --- |
| T0 | 120 | n.d. | 63 |
| T1 | 166 | 33 | 8 |
| T2 | 5 | 5 | 5 |

One hundred and sixty-six putative primary transformants (T0) were grown to maturity and tested. The first generation of seedlings (T1) were tested for resistance to hygromycin by germinating them on ½ MS medium supplemented with the antibiotics (Table 1). Only 33 lines germinated and grew, indicating that only 20 % were real transformants, of these, 8 lines (25 %) showed positive GUS staining after induction with Dex. Five of the most strongly inducible lines were grown to the next generation (T2), these lines retained HYG-resistance and showed positive GUS staining upon Dex induction. Two lines (65 and 121) were tested further and showed stable inducible GUS expression in the subsequent (T3) generation (Fig. 2A).

**Figure 2:**
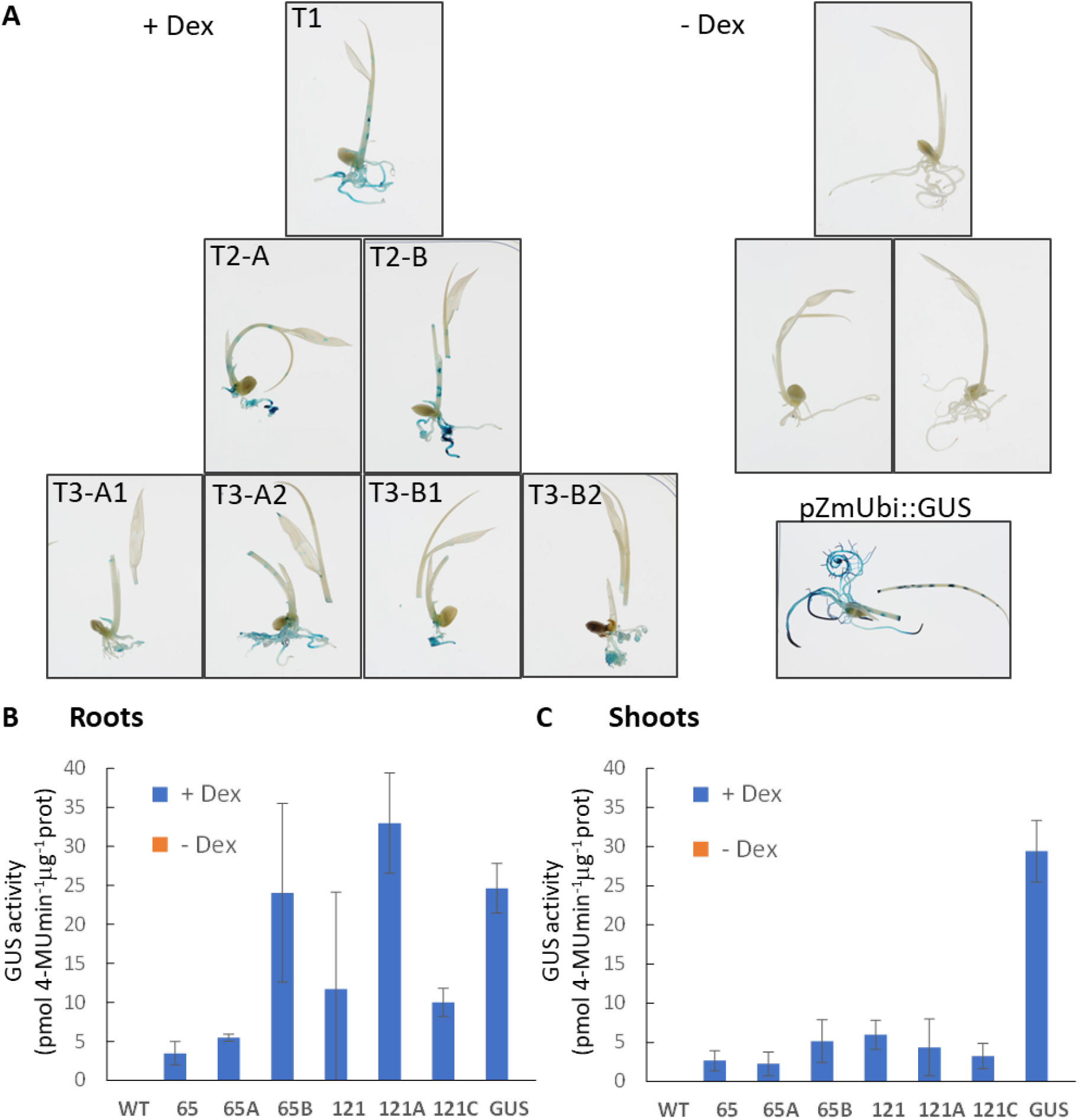
Reliability of inducible expression in subsequent generations and GUS activity in rice. **(A)** Ten-day-old rice seedlings (line 65) of generation T1, T2 (A and B) and T3 (A1, A2 and B1, B2) histochemically stained for GUS activity induced by germinating and growing the seedlings on ½ MS plates containing 30 μM Dex or control DMSO (-Dex) for 10 days. Seedlings of pZmUbi::GUS were included as a control for the GUS staining. **(B, C)** GUS activity determined fluorometrically in roots (B) and shoots (C) of 7-day-old rice seedlings (lines 65 and 121, T1 and T2) germinated and grown on ½ MS plates containing 30 μM Dex or control DMSO (-Dex) for 7 days. Seedlings of WT and pZmUbi::GUS were included as controls. The error bars represent SD.

We determined the GUS activity fluorometrically in segregating T1 and T2 progeny in roots (Fig. 2B) and shoots (Fig. 2C) of 7-day-old rice seedlings germinated and grown on ½ MS plates containing 30 μM Dex or the same concentration of DMSO (-Dex control). Interestingly, the induced GUS activity was up to 8-fold higher in roots compared to shoots and in some cases in roots it was comparable to the activity from the constitutive *pZmUbi* promoter. Perhaps a low transpiration rate in Petri dishes impaired the uptake and distribution of Dex into the shoots.

### Time course and dose response characteristics of Dex-induced GUS activity

To characterise the induction property of the pOp6/LhGR system in rice we performed time course and dose response experiments. To increase the efficiency of induction in shoots, we induced the newly developed leaves of app. 2-week old seedlings, grown in the open air, by painting the leaves with a Dex solution supplemented with 0.1% (v/v) Tween-20. Significant increase in fluorometrically, determined GUS activity, was detected 12h after induction with 10 μM Dex in two independent transgenic lines (65B and 121C) and this activity reached app. a half of the *pZmUbi* constitutive promoter activity within 72h of induction (Fig. 3A). The GUS activity was induced in plants treated with 0.01 μM Dex and while one line (65B) reached maximum levels of induction with 0.1 μM Dex, the other line (121C) had increased activity with increasing Dex concentration and reached levels similar to the constitutive *pZmUbi* promoter with 10 μM Dex after 48h induction (Fig. 3B).

**Figure 3:**
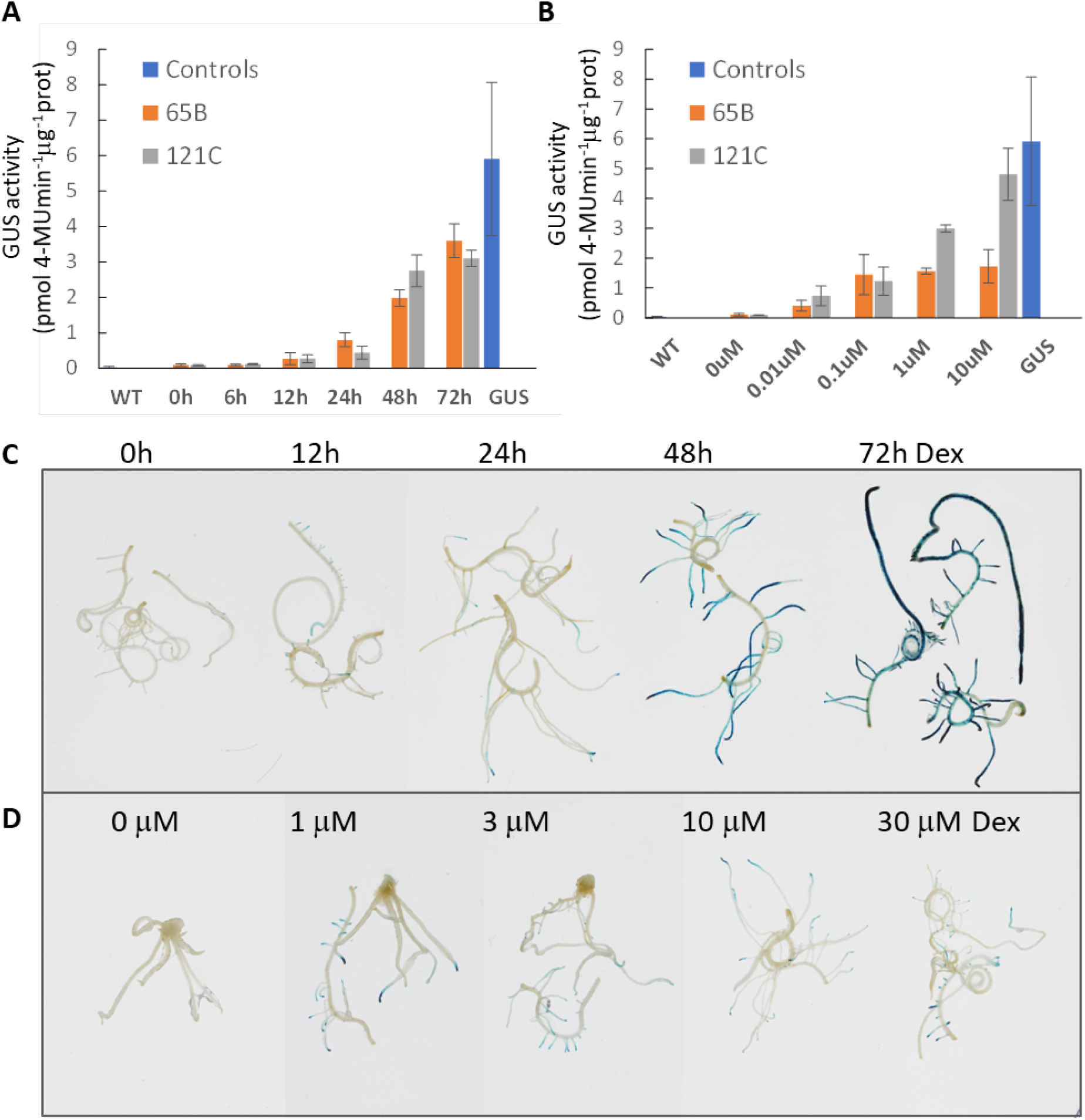
Time course accumulation of induced GUS activity and dose-response of the pOp6/LhGR system in rice. GUS activity determined fluorometrically in 7-day-old rice seedlings (lines 65B and 121C) germinated on ½ MS plates (vertically placed in the incubator) and then stood up in 15 ml Falcon tubes with 2 ml water for further 5 days. The newly developed leaves were induced by painting with water supplemented with 0.1% Tween-20 and **(A)** 10 μM Dex for 0, 6, 12, 24, 48 and 72h or **(B)** with 0, 0.01, 0.1, 1 and 10 μM Dex for 48h. Seedlings of WT and pZmUbi::GUS were included as controls. The error bars represent SD. **(C)** Roots of 10-day-old seedlings (lines 65A, 121C) were induced in liquid ½ MS with 30 μM Dex for 0, 12, 24. 48 and 72h, then histochemically stained for GUS activity. Representative images are shown. **(D)** Roots of 10-day-old seedlings (lines 65A, 121C) were induced in liquid ½ MS for 24h with 0, 1, 3, 10 and 30 μM Dex, then histochemically stained for GUS activity. Representative images are shown.

To confirm the similar characteristic of the pOp6/LhGR system in roots, we induced detached roots of 10-day old seedlings in liquid ½ MS media supplemented with increasing concentrations of Dex and performed histochemical GUS staining after specific time durations. Visible GUS staining was first detected in developing lateral roots and tips after 12h of induction and the intensity increased throughout the root system up to 72h tested (Fig. 3C). The maximal GUS staining intensity was detected with 1 μM Dex, the lowest concentration tested (Fig. 3D), predominantly in growing root tips after 24h induction.

We also tested the feasibility of inducing whole seedlings (10-day old) in a liquid ½ MS medium supplemented with 10 μM Dex. Histochemical GUS staining revealed the induction after 12h predominantly in roots and the staining pattern did not change significantly in the 48h time-span tested (data not shown).

### Optimization of induction by testing different steroids as inducers

We tried to improve the levels of induction of the pOp6/LhGR system in rice by testing different glucocorticoid derivatives (steroids) as inducers. In an attempt to reduce the surface tension at the air–liquid interface that is high in rice leaves due to epicuticular waxes preventing water vapor loss, we tested different concentrations of Tween-20 as the wetting agent rather than Silwet L-77 used previously (Craft *et al*., 2005; Samalova *et al*., 2005). Fig. 4A (i-iv) shows clear differences in the intensity of GUS staining of the 10-day-old shoots (leaves and stems) induced with a 30 μM Dex solution supplemented with 0.1% Tween-20 compared to 0.01%. Almost no staining was visible without the addition of the surfactant apart from damaged cells.

**Figure 4:**
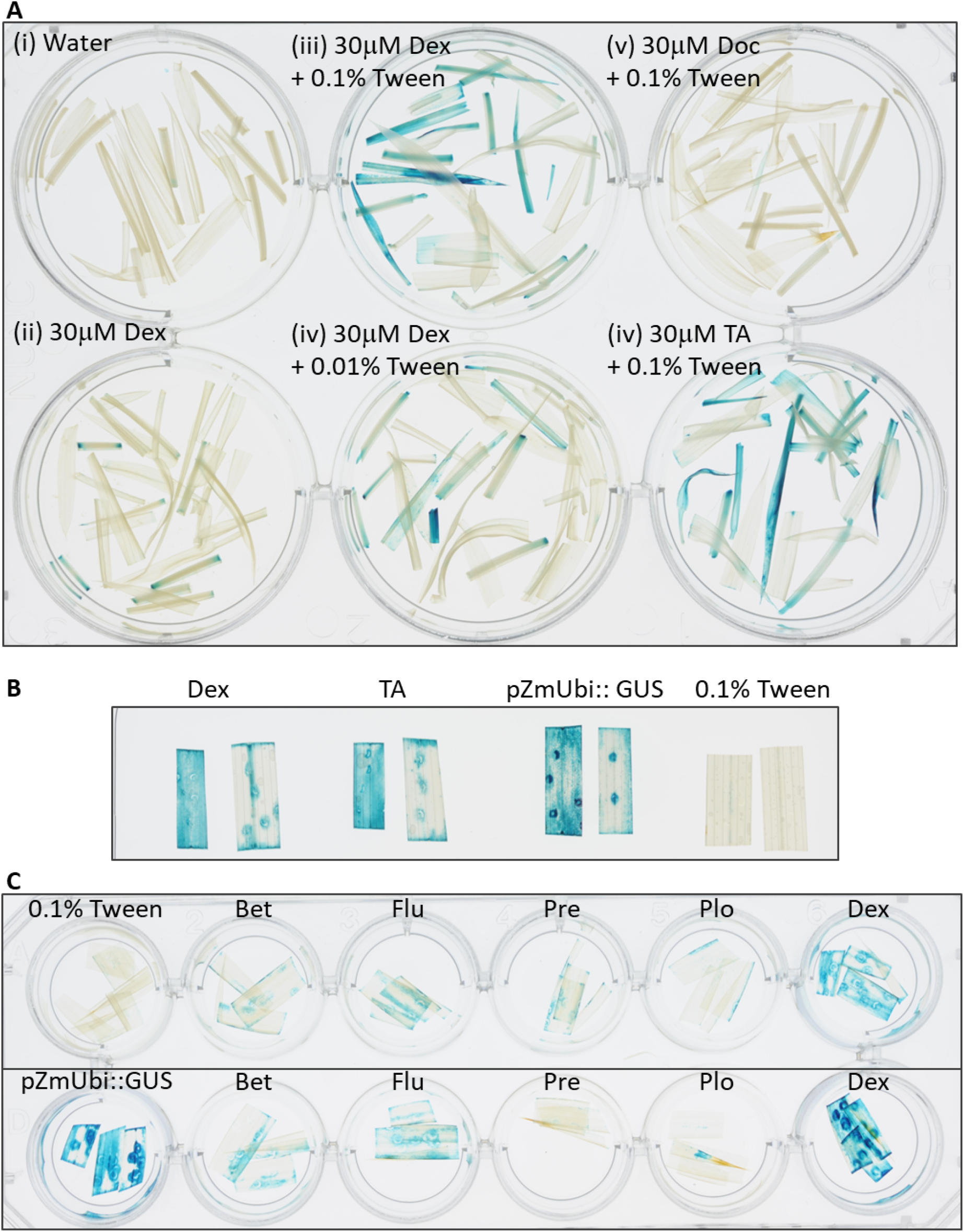
Optimization of the pOp6/LhGR induction in rice by testing different steroid inducers. **(A)** Shoots of 10-day-old seedlings (lines 65A, 121C) were induced in water supplemented with either 0.01% or 0.1% Tween-20 and 30 μM steroid inducer Dex, deoxycorticosterone (Doc) or triamcinolone acetonide (TA) for 24 h, then histochemically stained for GUS activity. **(B)** Leaf cuttings of the same leaf of 6-week-old plants (lines 65A, 121C) were induced in water supplemented with 0.1% Tween-20 and 10 μM Dex or 10 μM TA for 24h, then histochemically stained for GUS activity. Leaves of pZmUbi::GUS were included as controls. Representative images are shown. **(C)** Leaves of 5-week-old plants (lines 65A top, 121C bottom) were induced by painting with water supplemented with 0.1% Tween-20 and 30 μM steroid inducer: Dex, betamethasone (Bet), fludrocortisone acetate (Flu), prednisone (Pre) or prednisolone (Plo) for 24 h, then histochemically stained for GUS activity.

It is known that other glucocorticoid derivatives such as triamcinolone acetonide (TA) or deoxycorticosterone (Doc) can be used as inducers to replace Dex (Aoyama and Chua, 1997). We tested both in a 30 μM water solution supplemented with 0.1% Tween-20 (Fig. 4A v and vi) and compared the GUS staining intensities. Doc induction was negligible; however, TA induction was comparable if not higher than with Dex. To confirm this observation, we repeated the experiment on cuttings of the same leaf of 6-week-old plants that were submerged in water supplemented with 0.1% Tween-20 and 10 μM Dex or 10 μM TA for 24h (Fig. 4B). Depending on the efficiency of the substrate penetration, the GUS staining pattern with both inducers was comparable, but more importantly, also comparable to the staining of pZmUbi::GUS line of the same age. No GUS staining was detected without the inducers (Fig. 4B, 0.1% Tween-20).

We expanded the range of possible steroid inducers readily available and tested them in a 30 μM water solution supplemented with 0.1% Tween-20. Leaves of 5-week-old plants (Fig. 4C, lines 65A top and 121C bottom) were painted with the following steroids: betamethasone (Bet), fludrocortisone acetate (Flu), prednisone (Pre) or prednisolone (Plo) and Dex and histochemical GUS staining was carried out 24h later. None of the new glucocorticoids reached GUS activity levels comparable to Dex in both the transgenic lines tested and the GUS staining was considerably weaker than that of the constitutive *pZmUbi* promotor activity.

### Systemic and localised induction of soil-grown plants

To test the feasibility of inducing gene expression at later stages of plant development, either in a systemic or in a localized manner, Dex or the glucocorticoid TA were applied to soil-grown plants by watering (subterranean irrigation) or painting. Local induction of expression was detected after 24 h when leaves were painted with a 10 μM steroid solution supplemented with 0.1 % Tween-20 (Fig. 5A). The treatment was repeated and GUS activity determined 24 h and 72 h following first application of the inducer. Interestingly, the TA treatment doubled the induced levels of GUS activity compared to Dex, and both treatments exceeded the levels of *pZmUbi* promoter activity at 72 h post induction (hpi).

**Figure 5:**
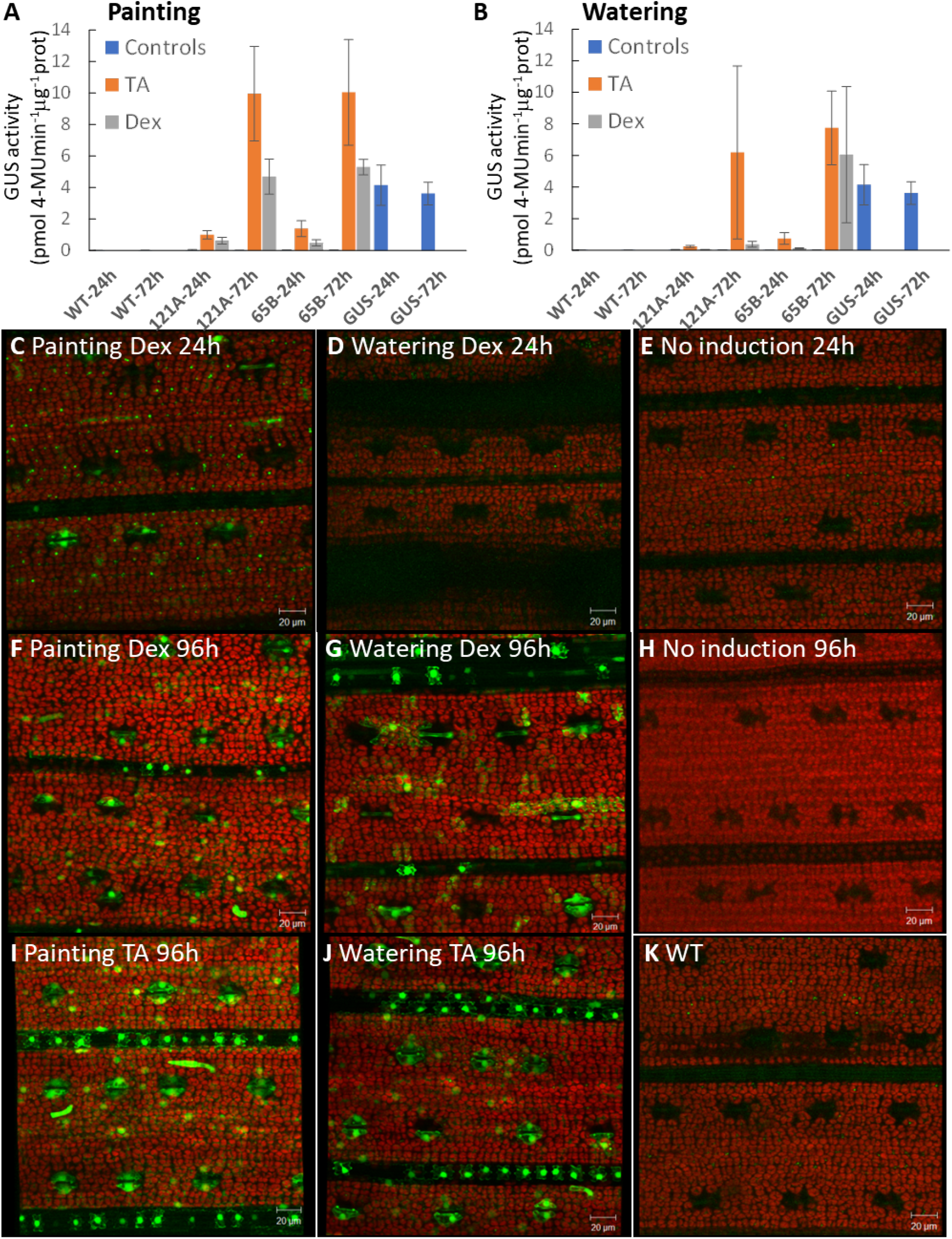
Expression characteristics of the pOp6/LhGR system induced in soil-grown plants by painting and watering. Four-week-old rice plants grown in soil were induced either with TA or Dex by **(A)** painting with 10 μM solution supplemented with 0.1 % Tween-20 or **(B)** watering with 30 μM solutions. The treatment was repeated twice at 0h and 24h later. Leaves were sampled and GUS activity determined 24h and 72h following first application of the inducer. WT and pZmUbi::GUS plants were used as controls. Three plants from segregating populations were used for each treatment. The error bars represent SD. **(C-K)** Four-week-old plants (line 121C) were induced as above except the treatment was repeated 48h after first induction. Leaves were imaged using a confocal laser scanning microscope at 24h and 96h post-induction, YFP signal in green, chlorophyll autofluorescence in red. Scale bar is 20μm.

Similarly, plants were watered with a 30 μM steroid solution and the treatment was repeated twice at 0h and 24h later. The induction triggered the reporter gene expression at the whole plant level, GUS activity was detectable in leaves 24 h after application of TA and 72h with both Dex and TA (Fig. 5B). The induced levels again exceeded the GUS activity detected in the pZmUbi::GUS line, however, but only with one of the lines tested (65B). The variability in the measurements could be due to segregating plant populations and a small sample size.

Both painting and watering experiments were repeated with a small modification of the treatment done at 2-day intervals and leaves were imaged using a confocal laser scanning microscope to observe induction of the second reporter gene, YFP (Fig. 5C-K). A bright YFP signal was detected 24h after painting the leaves (Fig. 5C) but not watering the plants (Fig. 5D) with Dex. The signal became stronger at 96 hpi with both methods of treatment (Fig. 5F and 5G) and more cells seemed to be expressing YFP with the TA treatment than Dex (Fig. 5I and 5J). No signal was detected without any induction (Fig. 5E and 5H) as in the WT (Fig. 5K).

To summarise, higher levels of GUS activity and *YFP* expression were obtained by painting the leaves rather than watering the plants and using TA at equivalent concentrations of Dex, suggesting that TA is a more potent inducer. Thus, whole plant and single leaf phenotypes can be assessed after induction using both methods.

### Long-term induction of the pOp6/LhGR system has no negative effects on rice plants

Final experiments were to determine whether there are any undesirable effects due to long term induction of the pOp6/LhGR system in rice. Four-week old plants were watered with either 30 μM Dex or 30 μM TA or a control solution (DMSO) at 2-day intervals for a week (Fig. 6, left) or two weeks and let to recover for further 6 weeks (Fig. 6, right). The images suggest that the plants grew and developed normally compared to WT plants treated with the control solution.

**Figure 6:**
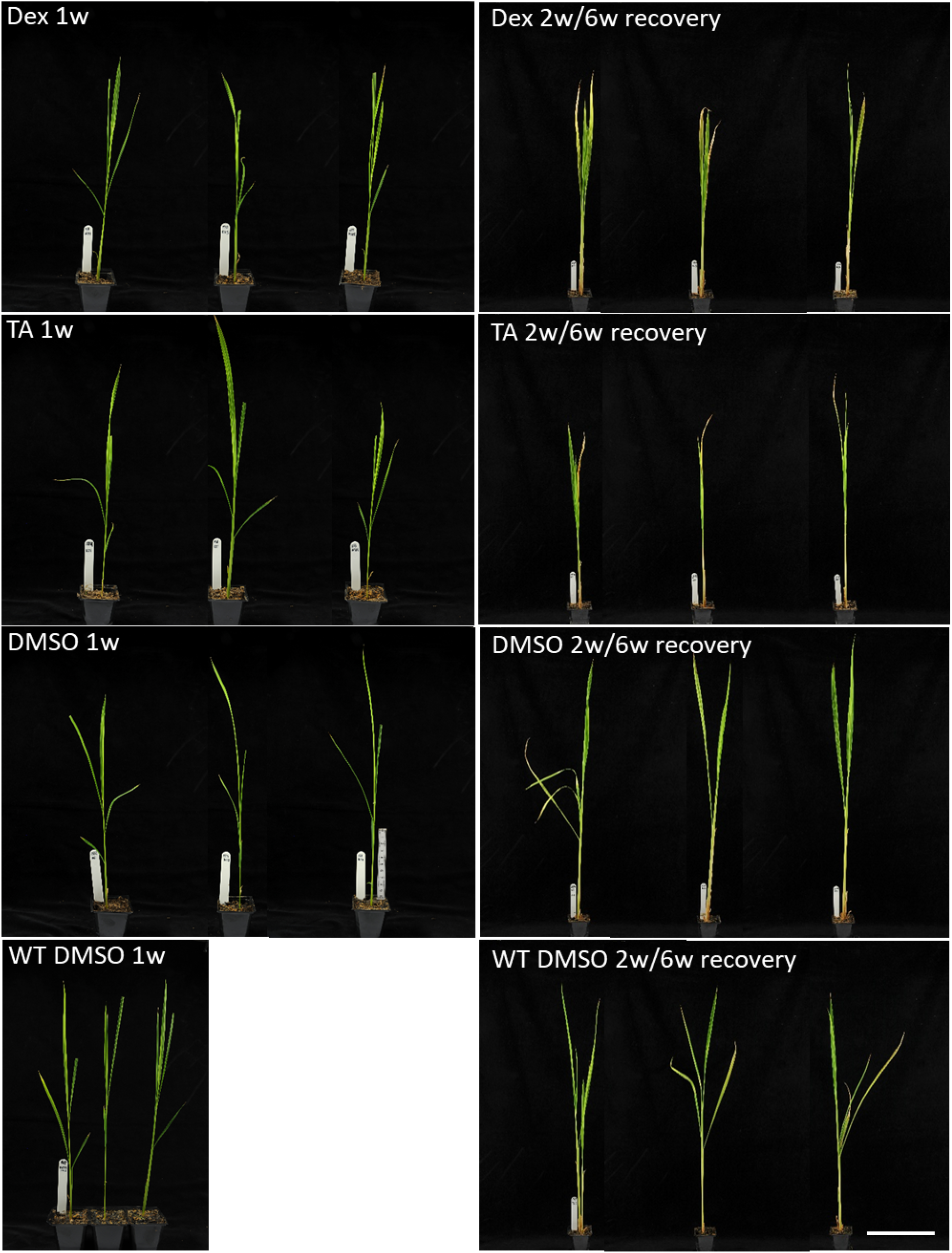
Long-term effect of the pOp6/LhGR system induction on mature rice plants. Four-week-old plants (65B and 121A) were watered with 30 μM Dex or 30 μM TA or a control solution containing DMSO (600 ml per a tray of 8 plants) at 2-day intervals for a week (left) or two weeks and let to recover for further 6 weeks (right). WT plants treated with the DMSO solution were used as controls. Scale bar represents 10 cm. Representative images are shown.

## DISCUSSION

### A construct design: possible reasons for toxicity, leakiness and silencing

In the publication of Vlad and colleagues (2019) the activator LhGR was codon optimized for use in rice (*rcoLhGR*) and further “domesticated” to remove all recognition sites for the Golden Gate cloning. However, severe growth defects on 10 μM Dex media were observed with all their constructs tested and in all transgenic lines. As addition of isopropyl β-D-1-thiogalactopyranoside (IPTG) to the growth media reduced the severity of the growth arrest phenotype, the authors concluded that it was a direct consequence of LhGR activity. However, in our work, we used the activator LhGR2 version that was only partially codon optimized for *Arabidopsis* to eliminate premature polyadenylation events in the Gal4 domain (Rutherford *et al*., 2005) and we did not observe any undesirable growth defects. Therefore, it is possible that the rice codon optimization rendered the system too efficient and prone to non-specific off-target binding due to high rcoLhGR protein levels in the growth inhibited lines. In this regard, to optimize efficiency of an expression system it is recommended to maximize the translational efficiency of a gene of interest (Moore *et al*., 2006). This can be achieved by taking some simple steps such as ensuring that the initiation codon conforms to the consensus for efficient initiation in plants (Koziel *et al*., 1996) and including translation enhancers in the 5’UTR such as the *TMV* Ω in our study.

Similarly, the GVG synthetic transcription factor that incorporates the glucocorticoid receptor (GR) ligand binding domain was found to be detrimental when activated in rice (Owerkverk *et al*., 2001) and other species including *Arabidopsis* (Kang *et al*., 1999; tobacco, Amirsadeghi *et al*., 2009; *Lotus japonicus*, Andersen *et al*., 2003). The growth perturbations were caused probably by the GVG activator that was binding to cis-regulatory elements in the plant genome with sequence homology to *GAL4* upon Dex activation (Kang *et al*., 1999). Nevertheless, the system was used successfully in a number of studies in rice (Chern *et al*., 2013; Nakayama *et al*., 2013; Onodera *et a*l., 2018, Park *et al*., 2012; Qu *et al*., 2009) in which the negative effects were reduced by shortening the time of Dex exposure or selecting lines with low GVG expression levels and mild phenotype that could be used as activator lines.

Moreover, when designing a construct for an inducible system, care must be taken to ensure that enhancers in the promoter that drives the activator do not activate the target promoter (*e*.*g*. pOp6; Moore *et al*., 2006) as such arrangement might result in transgene activation in the absence of inducer. In the pOp6/rcoLhGR constructs (Vlad *et al*., 2019), the arrangement was not ideal, as the pOp6 promoter was positioned directly next to the *pZmUbi* promoter driving the *rcoLhGR*. Furthermore, two other potent promoters were used on the same T-DNA; the constitutive rice actin promoter (*pOsACT*) to drive dsRed for identification of transgenic seeds, and another copy of *pOsACT* or the CaMV 35S promoter (for hygromycin-resistance selectable marker) that can also affect the expression of a transgene (Yoo *et al*., 2005). Curiously, the authors also used an intron in the reporter gene (GUS) placed again right next to the *pOp6* promoter and revealed that the presence of the intron had an enhancing effect on GUS activity levels both in the absence and presence of inducer.

Also, it must be recognized that any transgene may be susceptible to post-transcriptional gene silencing (PTGS) if transcripts accumulate to sufficient levels whether it is expressed constitutively or when induced (Moore *et al*., 2006). Individual reporter genes have a gene-specific threshold of mRNA abundance that will trigger PTGS (Schubert *et al*., 2004) and the probability of silencing of an inducible transgene locus in increased if the locus is induced (Abranches *et al*., 2005). In an attempt to minimize gene silencing, in our construct design we tried not to repeat the same sequence more than once, for example we used different polyadenylation signals (namely the octopine synthase terminator Tocs, nopaline synthase terminator Tnos, and T35S; Fig. 1A). However, we were not able to avoid silencing as only 25 % of hygromycin-resistant transgenic lines (T1) displayed GUS activity after Dex induction compared to 50 % of tested transgenic calli (T0). Others described a similar decrease, for example a reporter gene (GUSPlus) was expressed in 84% of rice calli but only in 25-68% of adult plants (Wu *et al*., 2003); and with the XVE system only 50 % of calli derived from inducible T1 seeds showed detectable GFP signal (Okuzaki *et al*., 2011).

### pOp6/LhGR compared with other systems: is there an ideal inducible system for rice?

The development of chemical-inducible systems for tight control of plant gene expression is a challenging task. There are a number of properties that are required for an ideal system, such as low basal expression level, high inducibility, specificity and dynamic range of response with respect to the inducer. Also, fast response and induction by various methods is desirable. An ideal system should work in several plant species and should not cause any adverse physiological effects in plants by itself or its inducer. The inducer is further required to show high specificity for the transgene, high efficiency at low concentrations and must not be found in target plants.

The pOp6/LhGR system described here **is fast**, first GUS activity was detected in painted leaves (Fig. 3A) and visible GUS staining in growing root tips (Fig. 3C) after 12h of induction and increased up to 72h tested. Similarly, with the rcoLhGR activator, 6h of induction was sufficient to induce high levels of GUS activity in transgenic calli and reached maximum levels at 24-96h depending on the construct (maximum levels of activity were reached after 24h in lines containing constructs with introns in the reporter gene but required 4 days of induction in the intron-less versions) (Vlad *et al*., 2019). On the other hand, the GVG system tested in rice (Ouwerkerk *et al*., 2001) required 4 days for the activity to be detected and 2 weeks of induction for GUS activities to reach levels comparable to those conferred by the strong *CaMV 35S* promoter. With the XVE system, a GFP signal was detected in rice calli 48 h after induction (with 1 μM 17-β-estradiol) and reached maximum (with treatment at 25 μM) in 8 days. However, in calli and leaves of transgenic lines the GFP signals were weaker than those in the leaves of p35S::GFP line, only in roots the signals were similar with 17-β-estradiol treatment at > 10 μM (Okuzaki *et al*., 2011).

The pOp6/LhGR system **is very sensitive**, 0.01 μM Dex was sufficient to induce GUS activity in painted leaves (Fig. 3B) and in roots of *in vitro* grown seedlings of the pOp6/rcoLhGR (Vlad *et al*., 2019). Interestingly, in some lines maximum levels of induction were reached with 0.1 μM Dex, while others required 10 μM Dex to reach levels similar to the constitutive *pZmUbi* promoter (Fig. 3B). This makes the pOp6/LhGR system ten-times more sensitive than the GVG system that required minimum of 0.1 μM Dex for induction and 1-10 μM for maximum.

The pOp6/LhGR system **is tightly regulated and strong**, the average fold induction was app. 500 in shoots of plants induced *in vitro* (Fig. 2C) or in soil by Dex, however, reached 1000-fold induction when plants were treated with TA (Fig. 5). In roots the magnitude of Dex-induction reached at least 1000-to 6000-fold (Fig. 2B), similar to the values reported for the GVG system (Ouwerkerk *et al*., 2001). Both the pOp6/LhGR as well as the pOp6/rcoLhGR systems used the same line that constitutively expressed a synthetic GUS variant from *Staphylococcus* (GUSPlus) under the control of the *pZmUbi* promoter as a positive control. The GUSPlus was reported to be ten-fold more active than the enzyme encoded by *E. coli uidA* that was used in the inducible constructs (Broothaerts *et al*., 2005; Jefferson *et al*., 2003), suggesting that activation of the pOp6 promoter by the activators is very effective.

The pOp6/LhGR system **is inducible by various methods** *in vitro* and in soil-grown plants. Painting the leaves with an inducer solution supplemented with 0.1% Tween-20 (to help the inducer to penetrate through the protective wax layer) proved to be more effective than subterranean irrigation (watering). It is worth to note that using these methods of application, TA proved to be almost twice as more an efficient inducer than Dex (Fig. 5). This could be due to the nature of the inducer as Dex could be more rapidly metabolized or compartmentalized than TA (Aoyama and Chua, 1997). Ouwerkerk and colleagues (2001) reported that Dex was taken up efficiently by roots of mature plants in hydroponics and induced GUS activity throughout the whole plant body.

The major limitation and possible reasons for low levels of target gene induction in rice leaves using the XVE system is the inefficient uptake of estradiol from hydroponics. Interestingly, cut leaf segments placed in liquid MS media with 10 μM estradiol showed detectable GFP fluorescence in whole leaf segments within 48 h (Okuzaki *et al*., 2011) and similar activation was confirmed by PCR in another study (Hirose *et al*., 2012). Also, spraying of leaves with the inducer was another effective method of induction (Hirose *et al*., 2012). The limiting factor therefore seems to be the estradiol uptake that is depending on the roots. The inducer might have been trapped by XVE expressed in roots or diffused and attenuated in leaves. Interestingly, activation by soaking transgenic seeds (Chen *et al*., 2017) with estradiol solution induced highly efficient site-specific recombination in germinating embryos, resulting in high-level expression of target gene or RNAi cassette in intact rice plants.

The pOp6/LhGR system **is not toxic in rice** and does not cause any undesirable negative developmental growth effects even at higher concentration of Dex (up to 30 μM tested) and after prolonged exposure of the plants (Fig. 6). However, for the long-term induction through soil, DMSO should be avoided and ethanol should be used as a solvent for the inducers to prevent accumulation of DMSO in the soil over several weeks of watering (Moore *et al*., 2006; Samalova *et al*., 2019). Very unexpected were the findings of Vlad and colleagues (2019) that the pOp6/rcoLhGR transgenic plants manifested severe developmental perturbations when grown on concentrations > 0.1 μM Dex. The direct cause of these growth defects is not known, but the authors suggested that the rice genome contains sequences with high similarity to the *lacOp* sequence, suggesting non-specific activation of endogenous genes by Dex induction. Although, the pOp promoter was not activated by endogenous factors in maize (Moore *et al*., 2006; Segal *et al*., 2003) it is possible that the rice-codon optimization rendered the system too efficient. The off-target effects can be minimized by quenching with IPTG that acts as an effective antagonist for the LacI DNA binding domain in LhGR (Craft *et al*., 2005).

The GVG system has been shown to generate growth defects in rice (Owerkverk *et al*., 2001) and other species (Kang *et al*., 1999; Andersen *et al*., 2003; Amirsadeghi *et al*., 2009) but the reason for this is not clear as each domain of the molecule has been used successfully in other expression systems. However, one possibility is that GVG requires higher expression levels to achieve full promoter activation, resulting perhaps in squelching or non-specific binding to *CCG(N11)CGG* sequences in the genome (Moore *et al*., 2006). Apparently, the GVG level was not limiting for target-gene activation (Aoyama and Chua, 1997), therefore, selection of plants with a mild GVG phenotype in combination with optimized induction conditions was recommended for the system applications. Although, the XVE system had no effect on growth, all tested lines including controls grown in hydroponics presented a different root pattern with oestradiol treatment so the system does not appear to be suitable for root morphology studies (Okuzaki *et al*., 2011).

## EXPERIMENTAL PROCEDURES

### Construct preparation

To prepare the pVecLhGR2 construct (Fig. 1A, Fig. S1), first, the LhGR2 and the octopine synthase terminator (Tocs) sequences were amplified by polymerase chain reaction (PCR) using vector pOpIn2 (Samalova *et al*., 2019) as the template. The two PCR fragments, LhGR2 (forward primer P1: 5’-AAAAAGGTACCATGGCTAGTGAAGCTCGAAAAACAAAG-3’; reverse primer P2: 5’-GCATATCTCATTAAAGCAGGACTCTAGTTCACTCCTTCTTAGGGTTAGGTGGAGTATC-3’) and Tocs (forward primer P3: 5’-TACTCCACCTAACCCTAAGAAGGAGTGAACTAGAGTCCTGCTTTAATGAGATATGC-3’; reverse primer P4: 5’-AAAAAGGTACCCTAAGGCGCGCCGTTGTCG CAAAATTCGCCCTGGACCC -3’) were joined together by overlapping PCR with primers P1 and P4 and cloned into the unique *Kpn*I site of the binary rice overexpression vector pVec8-*Kpn*IOX (Kim and Dolan, 2016). Secondly, an *Asc*I fragment containing the bidirectional pOp6 operator array together with *TMV* Ω translation enhancer sequences driving expression of YFP and GUS (including 35S terminator sequence) reporters of the pOpIn2YFP vector (generated by M. Kalde, I. Moore lab) was cloned into a unique *Asc*I site created in the previous cloning step. The correct orientation of the fragment as well as the LhGR2 in the final pVecLhGR2 vector was confirmed by sequencing.

### Plant material and growth conditions

To generate stable transgenic rice lines, we used *Agrobacterium-*mediated transformation of *Oryza sativa spp. japonica* cultivar Kitaake. Calli were induced from dehulled mature seeds before co-cultivation with *A. tumefaciens* strain EHA105 transformed with the pVecLhGR2 construct. Callus transformation and seedling regeneration were performed at 28 °C according to a protocol modified from Toki *et al*., (2006) as described by Vlad *et al*., 2019. Independent pVecLhGR2 lines were tested together with positive *pZmUbi*::*GUS* (Vlad *et al*., 2019) and negative wild-type (WT) controls. Soil-grown plants were cultivated in a greenhouse at 28-30 °C with a 16h/8h photoperiod.

### Methods of steroid induction – *in vitro*, painting and watering

Dexamethasone (Dex) was prepared as a 30 mM stock solution in dimethyl sulfoxide (DMSO) and stored at −20 °C. Similarly, we prepared stock solutions of betamethasone (Bet), deoxycorticosterone (Doc), fludrocortisone acetate (Flu), prednisone (Pre), prednisolone (Plo) and triamcinolone acetonide (TA), all purchased from Sigma-Aldrich. For each treatment, either a glucocorticoid (induction) or the equivalent volume of DMSO (control) were added to obtain desired concentration.

For application *in vitro*, seedlings were grown on half strength Murashige and Skoog (1962) medium (½ MS medium) supplemented with 15g/l sucrose and with or without agar. Dex was added to the medium after autoclaving to obtain final working concentration, typically 30 μM Dex. Shoots and leaf cuttings were also induced in distilled water supplemented with 0.1% Tween-20 and a glucocorticoid inducer (10-30 μM).

For induction of soil grown plants, 4-week-old rice plants were either watered with 50 ml of 30 μM Dex or 30 μM TA or newly developed leaves (3^rd^ leaf) were painted on both sides with 10 μM Dex or 10 μM TA solution supplemented with 0.1% Tween-20. The treatments were repeated at 2-day intervals for one or two weeks.

### Analysis of ß-glucuronidase (GUS) reporter activity

Histochemical GUS staining and fluorometric GUS assay was performed according to Jefferson (1987) at 37 °C. Extracts were prepared from ground snap-frozen samples; the protein content was determined spectrophotometrically using Bio-Rad Protein Assay (Bio-Rad laboratories). The fluorogenic reaction was carried out in 96 well-plates using protein extraction buffer supplemented with 4-methylumbelliferyl ß-D-glucuronide (4-MUG) as a substrate. A standard curve of 4-MU was used to calculate the amount of 4-MU/unit of time. Activity was calculated from three technical replicates and expressed in pmoles 4-MU/min /μg protein.

### Detection of YFP fluorescence

The YFP signal was detected using a Zeiss LSM 510 Meta confocal laser-scanning microscope with 514-nm excitation from an Argon laser and a BP535-590IR emission filter.

## ACKNOWLEDGEMENT

I would like to dedicate this publication to my colleague and friend Ian Moore, and his beloved family, without whom science is much less fun. I want to thank to Peng Wang (Chinese Academy of Sciences, Shanghai, China) for the control pZmUbi::GUS line and the rice transformation protocol; to Chul Min Kim (University of Oxford) for the pVec8-*Kpn*IOX vector and to Monika Kalde (University of Oxford) for the pOpIn2YFP vector. I am grateful to Molly Dewey for her suggested word changes.

## SUPPLEMENTARY FIGURE

**Figure S1:**
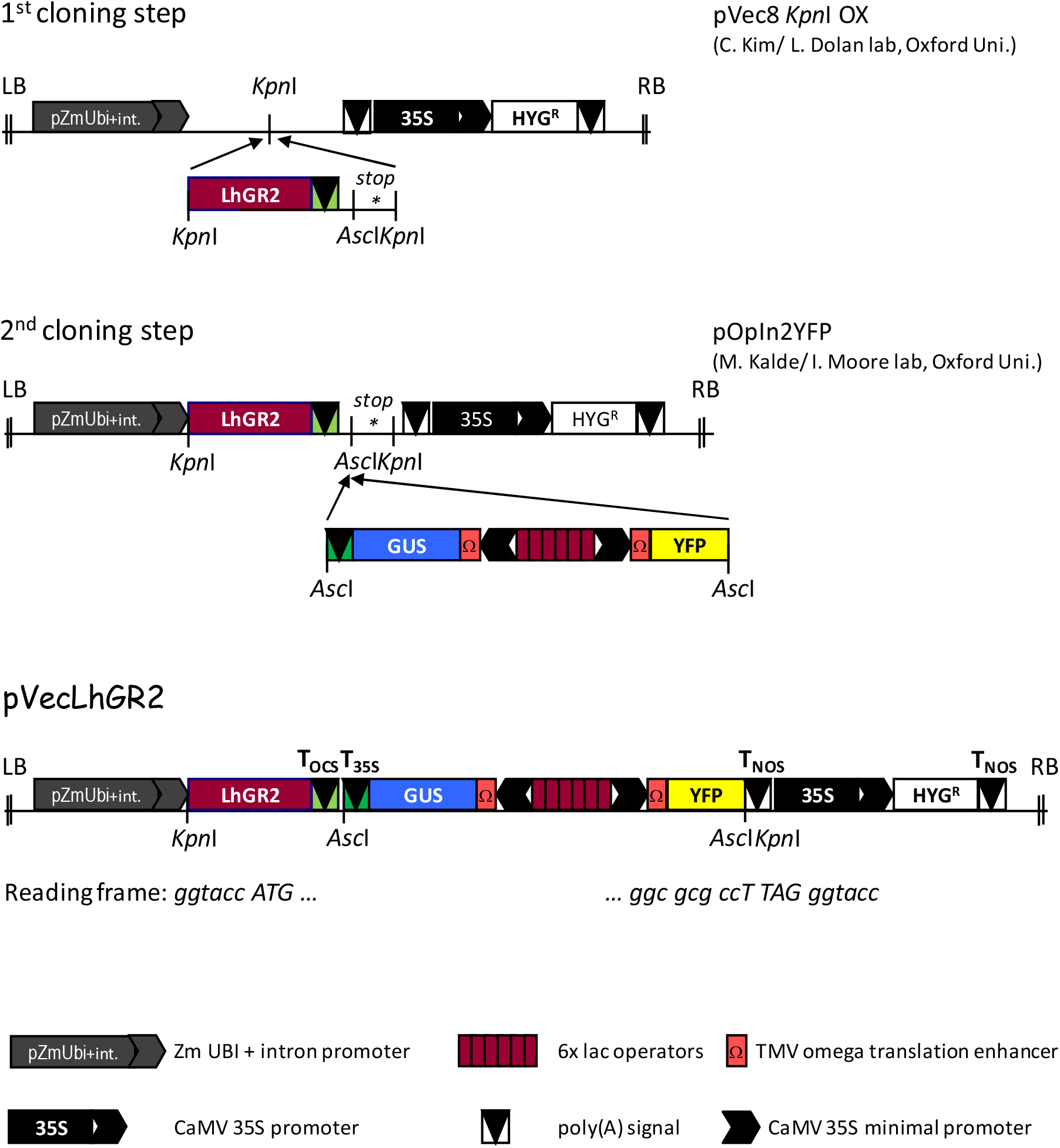
Cloning strategy for preparation of the pVecLhGR2 construct.

